# Distinct TORC1 signalling branches regulate Adc17 proteasome assembly chaperone expression

**DOI:** 10.1101/2023.12.11.571133

**Authors:** Thomas D. Williams, Sylwia M. Dublanska, Rebecka Bergquist, Adrien Rousseau

## Abstract

When stressed, cells need to adapt their proteome to maintain protein homeostasis. This requires increased proteasome assembly. Increased proteasome assembly is dependent on increased proteasome assembly chaperone expression. In *S. cerevisiae*, inhibition of the growth-promoting kinase complex TORC1 causes increased proteasome assembly chaperone expression. This is dependent upon activation of the MAPKinase Mpk1 and relocalisation of assembly chaperone mRNA to patches of dense actin. We show here that TORC1 inhibition alters cell wall properties to induce these changes by activating the Cell Wall Integrity pathway through the Wsc1, Wsc3, and Wsc4 sensor proteins. We demonstrate that in isolation these signals are insufficient to drive protein expression. We identify that the TORC1-activated S6Kinase Sch9 must be inhibited as well. This work expands our knowledge on the signalling pathways which regulate proteasome assembly chaperone expression.

**Summary:** TORC1-regulated proteasome assembly chaperone expression is necessary for proteasome assembly and proteostasis. We identify Cell Wall Integrity and Sch9 signalling as mechanisms controlling this.

## Introduction

When cells are stressed, they need to rapidly adapt their proteome to maintain protein homeostasis and survive (Williams and Rousseau, 2022). Proteome adaptation requires both an increase in the degradative capacity of the cell, either through increased autophagy or proteasome assembly, and changes to the proteins being produced through altered transcription, translation, or both. The environmental-sensing protein kinase complex TORC1 (mTORC1 in mammals) co-ordinates the processes of protein translation and degradation at a bulk level (Williams and Rousseau, 2022). When these interlocking aspects of protein homeostasis are perturbed, including during aging, diseases follow (Rousseau and Bertolotti, 2018).

Proteasome assembly induction upon nutrient stress is conserved from yeast to mammals (Gao et al., 2015; Rousseau and Bertolotti, 2016). The resulting increase in degradative capacity relies upon increased translation of proteasome assembly chaperones, including the yeast-specific stress inducible chaperone Adc17. Adc17 is commonly used as a model protein to explore increased proteasome assembly chaperone induction upon stress (Rousseau and Bertolotti, 2016; Williams et al., 2022; Williams, In press). The induction of Adc17 expression at the level of translation is now fairly well described: translation of *ADC17* mRNA is reliant upon Mpk1 activation and mRNA relocalisation to actin patches following actin depolarisation (Figure 1A) (Rousseau and Bertolotti, 2016; Williams et al., 2022; Williams, In press).

**Figure 1:**
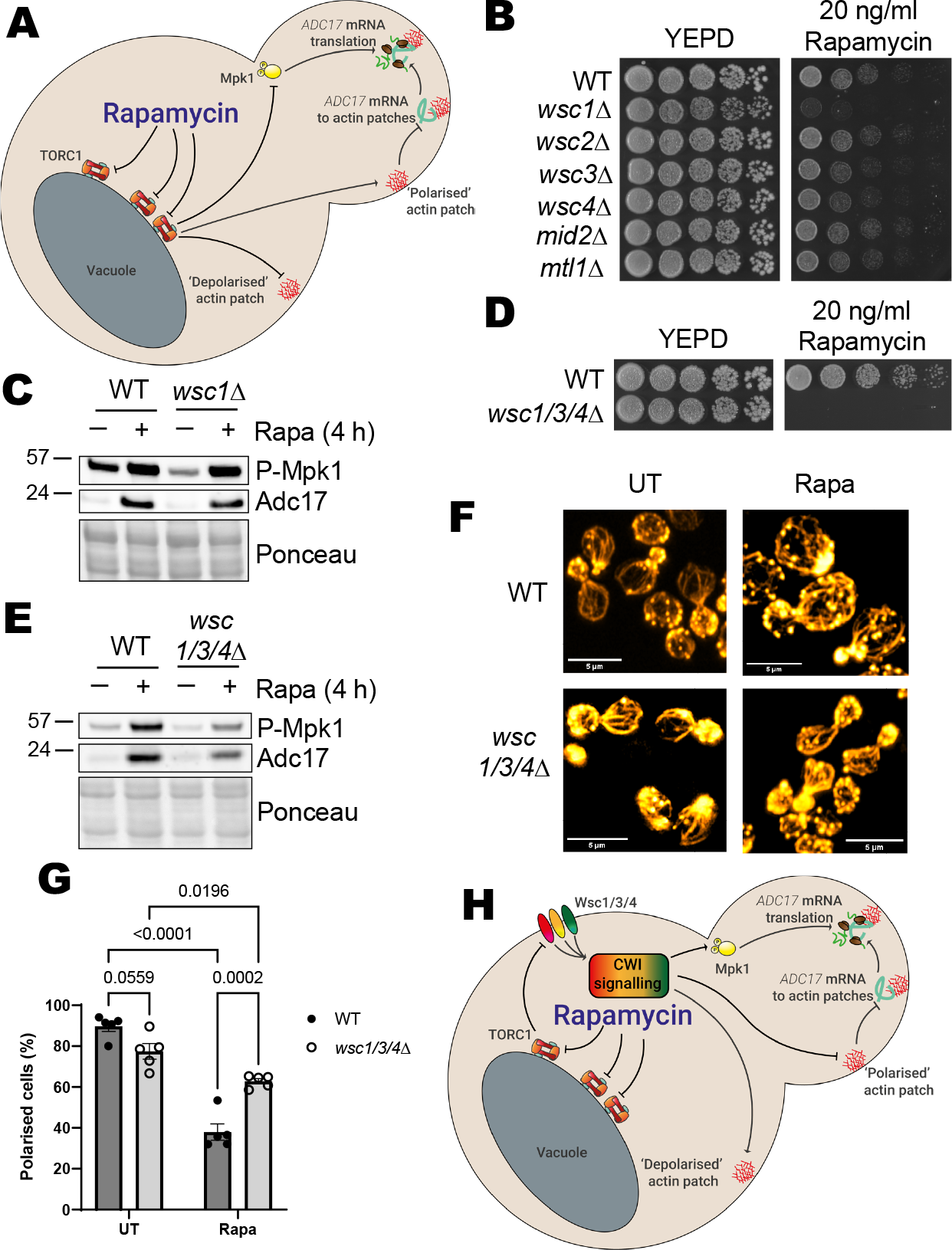
Mpk1 is activated and actin depolarised via the wsc1/3/4 cell wall integrity pathway sensors in response to rapamycin treatment. A) Proteasome assembly chaperone expression regulation overview. Upon rapamycin treatment, TORC1 is inhibited, leading to activation of Mpk1 and *ADC17* mRNA relocalisation to actin patches, together driving increased Adc17 expression. B) Drop assay of WT and CWI sensor mutants on YEPD and YEPD + rapamycin after 3 days of growth. Representative of multiple experiments. C) Mpk1 activation (P-Mpk1) and Adc17 expression following 4 h rapamycin treatment in WT and *wsc1Δ* cells. Representative of at least 3 experiments. D) Drop assay of WT and *wsc1/3/4Δ* cells on YEPD and YEPD + rapamycin after 3 days of growth. Representative of multiple experiments. E) Mpk1 activation (P-Mpk1) and Adc17 expression following 4 h rapamycin treatment in WT and *wsc1/3/4Δ* cells. Representative of at least 3 experiments. F) Effect of 1 h rapamycin treatment on actin polarisation in WT and *wsc1/3/4Δ* cells. Representative of at least 3 experiments. Cellular actin is labelled with rhodamine-phalloidin. Scale bars show 5 µm. G) Quantification of images from E (n=5). H) Updated scheme from Figure 1A, showing role of Wsc1/3/4 and CWI signalling in Adc17 expression.

Specifically inhibiting TORC1 via rapamycin treatment has long been known to activate Mpk1 and cause actin depolarisation (Krause and Gray, 2002; Torres et al., 2002). Both actin depolarisation and Mpk1 activation are well described as being downstream effects of Cell Wall Integrity (CWI) pathway activation (Jiménez-Gutiérrez et al., 2020). The CWI pathway was initially described as being activated in response to cell wall stress but has since been shown to be activated by many other stresses through the sensor proteins Wsc1, Wsc2, Wsc3, Wsc4, Mid2, and Mtl1. Wsc1 is the best characterised of these. Wsc1 has a cytosolic tail, a transmembrane domain, an extracellular ‘spring’ domain, and a cysteine rich head domain which is required for clustering in distinct microdomains, and possibly interaction with the cell wall (Dupres et al., 2009; Dupres et al., 2011; Kock et al., 2016; Schöppner et al., 2022). Perturbations affecting the cell wall cause a conformational change in the cytosolic domain, allowing interaction with and activation of the RhoGEF Rom2 (Jendretzki et al., 2011). Rom2 activates Rho1, which triggers a kinase activation cascade from Pkc1 to Mpk1 via Bck1 and Mkk1/2.

While Mpk1 is activated downstream of TORC1 inhibition or CWI activation, it remains unclear whether TORC1 inhibition activates the CWI pathway, or merely has a similar output. There have been conflicting reports on the potential role of the CWI sensor Mid2 in Mpk1 activation following TORC1 inhibition, while Wsc1 may also have a role (Krause and Gray, 2002; Torres et al., 2002). *wsc1Δ* and *mid2Δ* cells are not known to be rapamycin-intolerant, as would be expected with impaired Mpk1 activation (Rousseau and Bertolotti, 2016). It therefore remains an open question whether Mpk1 is activated primarily through the CWI pathway following rapamycin treatment. To further complicate matters, actin depolarisation can activate the CWI pathway, as well as being caused by CWI activation (Harrison et al., 2001). This raises the question of whether actin depolarisation induced by TORC1 inhibition is CWI-dependent or independent, in addition to Mpk1 activation.

Here, we clarify the links between TORC1, the CWI, actin architecture, and Mpk1 activation. We show that TORC1 inhibition indeed activates the CWI to induce Mpk1 activation and actin rearrangements, but that this is not sufficient for Adc17 expression. We identify that the S6Kinase Sch9 must also be inhibited for Adc17 expression. This work increases our knowledge of the signalling governing proteasome assembly chaperone expression.

## Results & Discussion

### TORC1 regulates the CWI sensor proteins Wsc1, Wsc3, and Wsc4 to control Mpk1 activation

To obtain more detail about how the proteasome assembly pathway is induced following rapamycin treatment, we began by investigating the mechanisms of Mpk1 activation. Cells lacking Mpk1 are incapable of growth in the presence of rapamycin, a phenotype which is also observed in the upstream kinase deletion mutants *bck1Δ* and *mkk1Δ/mkk2Δ* (Rousseau and Bertolotti, 2016). Therefore, strains that are incapable of Mpk1 activation following rapamycin treatment will likewise be incapable of growth in the presence of rapamycin. As Mpk1 is well described as being activated by a kinase cascade initiated by the Cell Wall Integrity (CWI) pathway, we focused on the most upstream element of this pathway: the CWI sensors.

We assessed the ability of knockouts for each of the 6 CWI sensor proteins to grow on rapamycin-containing plates (Figure 1B). All 6 mutants showed some level of growth, indicating functional redundancy between the sensor proteins. Three of the mutants, *wsc1Δ, wsc3Δ*, and *wsc4Δ* showed a mild to moderate growth defect. The most severely affected strain, *wsc1Δ*, has previously been implicated in Mpk1 activation following rapamycin treatment (Krause and Gray, 2002). We did not observe an Mpk1 activation defect using our *wsc1Δ* strain and conditions however (Figure 1C).

Reasoning that there may be functional redundancy between the Wsc1, Wsc3, and Wsc4 sensor proteins, we made a *wsc1/3/4Δ* strain. We found that *wsc1/3/4Δ* cells are incapable of growth in the presence of rapamycin (Figure 1D) and have greatly diminished Mpk1 activation following rapamycin treatment (Figure 1E). TORC1 inhibition therefore leads to activation of Mpk1 through the CWI pathway via the redundant activation of the CWI sensor proteins Wsc1, Wsc3, and Wsc4.

### CWI activation leads to actin depolarisation independently of Mpk1 activation

We next examined whether CWI activation caused by TORC1 inhibition leads to actin depolarisation, or whether the actin depolarisation leads to CWI activation. We analysed the effect of rapamycin on actin polarisation in *wsc1/3/4Δ* cells, where the CWI is impaired following rapamycin treatment (Figure 1E&F). Following rapamycin treatment, WT cells become largely depolarised, while the *wsc1/3/4Δ* cells remain significantly more polarised. Actin depolarisation therefore occurs downstream of CWI activation following TORC1 inhibition.

We tested whether actin depolarisation was regulated by Mpk1 activation. Deletion of Mpk1 did not inhibit actin depolarisation upon rapamycin treatment (Figure S1A&B), unlike previously observed in heat stressed cells (Guo et al., 2009). Activating Mpk1 by inducible expression of a constitutively active version of the upstream kinase Mkk1, Mkk1-DD (Harrison et al., 2004), likewise had no impact on actin polarity (Figure S1C-E). CWI regulation of actin polarity is therefore Mpk1-independent.

### CWI-Bni1-actin polarity section deleted as suggested, ready for the reviewers if required

These data suggest that TORC1 inhibition activates the CWI through the Wsc1, Wsc3, and Wsc4 sensor proteins. CWI activation then causes both actin depolarisation and Mpk1 activation to regulate proteasome assembly chaperone translation (Figure 1G).

### TORC1 inhibition activates the CWI by altering cell wall properties

It is unclear how inhibiting TORC1, a predominantly vacuole membrane-associated kinase complex, would activate Wsc1, Wsc3, and Wsc4 at other membranes. While other reports suggest a potential role for the TORC1-regulated phosphatase Sit4, elevated Mpk1 activity in *sit4Δ* cells makes this unlikely (Torres et al., 2002). Linking the cell wall and TORC1, previous work showed TORC1 inhibition affects cellular resistance to zymolyase, a yeast cell wall degrading enzyme (Krause and Gray, 2002). The observed increased zymolyase resistance, indicative of alterations in the cell wall, occurs after several hours of rapamycin treatment and is affected by the CWI-activated kinase Pkc1. When cells are provided with osmotic support (which can stabilise cells with weak cell walls), Mpk1 activation in response to rapamycin treatment is reduced (Torres et al., 2002). This suggests an effect at the cell wall may occur much earlier than previously described.

We assessed whether TORC1 inhibition affected cell wall properties by analysing zymolyase resistance in WT cells during the initial hour following rapamycin treatment (Figure 2A). A trend of increasing zymolyase resistance was apparent from 5 minutes post-treatment, although this didn’t become statistically significant until after 30–60 minutes. We examined whether this increased resistance was CWI-dependent using the *wsc1/3/4Δ* strain, where CWI activation is impaired (Figure 1E). WT and *wsc1/3/4Δ* cells which had not been treated for rapamycin had a comparably low resistance to zymolyase. For both strains 60 minutes of rapamycin treatment caused a comparable increase in resistance to zymolyase (Figure 2B). This indicates that there is no requirement for CWI activation for the initial cell wall changes responsible for increased zymolyase resistance. Increased resistance to zymolyase is therefore an early response to rapamycin treatment, before being later modulated by CWI activation.

**Figure 2:**
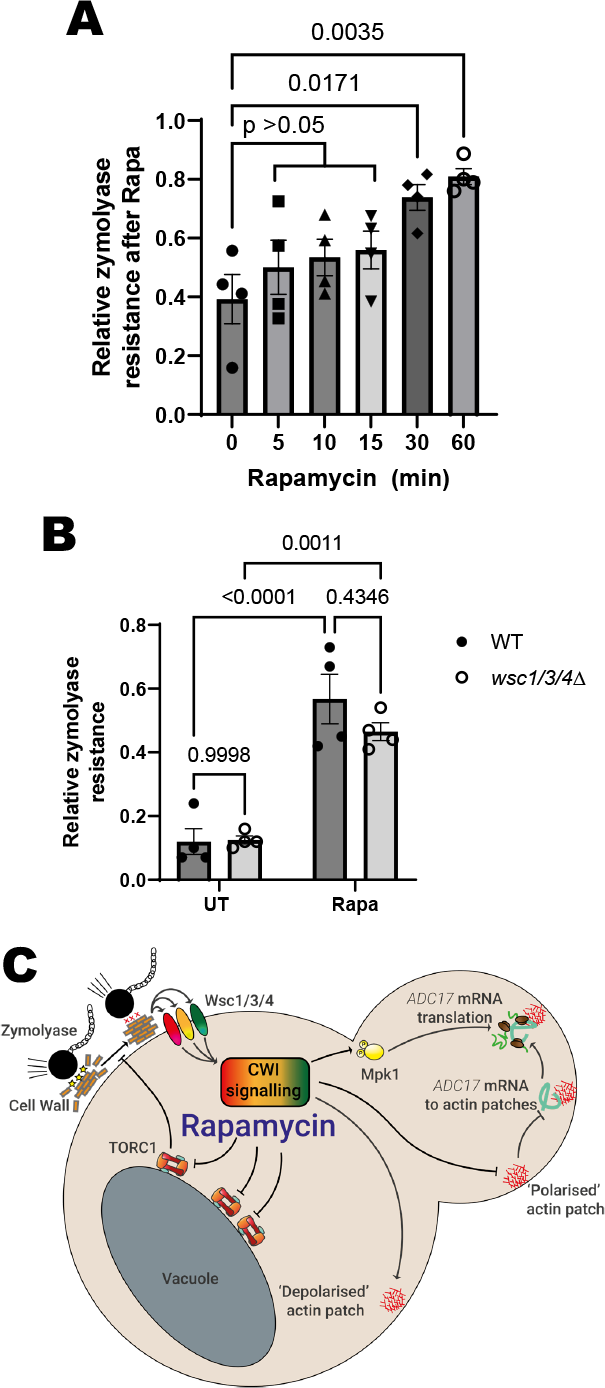
Rapamycin treatment activates cell wall integrity pathway by altering cell wall properties. A) WT cell resistance to 45 min zymolyase treatment after treatment with rapamycin (n=4). B) Resistance to 45 min zymolyase treatment by WT and *wsc1/3/4Δ* cells either untreated or after 1 h rapamycin treatment (n=4). C) Updated scheme from Figure 1A, showing role of cell wall alterations which increase zymolyase resistance in Adc17 expression.

TORC1 inhibition affects many downstream signalling branches, notably resulting in autophagy activation (Rabanal-Ruiz et al., 2017), S6Kinase inhibition (Urban et al., 2007), and PP2A phosphatase activation through Tap42 inhibition (Hughes Hallett et al., 2014). We used genetic manipulation to prevent these signalling changes from occurring (Figure S2A). If a particular branch was required for this change, there should be a decrease in the zymolyase resistance observed after rapamycin treatment. Individually preventing activation of autophagy, the Pph2-1/Pph2-2/Sit4 phosphates, or inhibition of the major yeast S6Kinase Sch9 had no effect on zymolyase resistance (Figure S2B).

The largest change was observed when a constitutively-active phospho-mimetic Sch9-2D3E expression construct was used (Urban et al., 2007). Reasoning that there may be functional redundancy, we expressed this construct in the other strains and again tested zymolyase resistance in response to rapamycin treatment. When prevention of Sch9 inhibition was combined with prevention of activation of any of the phosphatases (Pph2-1, Pph2-2, or Sit4) we observed a statistically significant, albeit rather small, decrease in the resistance to zymolyase (Figure S2C). In summary, TORC1 inhibition activates the CWI through altering cell wall properties to increase cellular zymolyase resistance (Figure 2C). The nature of these changes, and the signalling processes involved, are an open question to be explored in later studies.

### CWI activation induces Adc17 expression

To gain further detail on proteasome assembly pathway regulation, we examined whether rapamycin-induced CWI activation was sufficient for Adc17 expression. We therefore bypassed TORC1, activating the CWI pathway directly using the chitin-binding compound calcofluor white (CFW), which is well described as a CWI activator (Elorza et al., 1983). WT cells treated with CFW showed strong CWI induction (assessed by increased P-Mpk1 expression) and Adc17 expression (Figure S3A). Adc17 expression was dependent on Mpk1 (Figure S3A). We also observed the expected actin depolarisation and *ADC17* mRNA relocalisation to actin patches (Figure S3B-D). However, TORC1-dependent phosphorylation of Rps6 was reduced following CFW treatment (Figure S3E), likely through the action of Rho1 (Yan et al., 2012), preventing us from ruling out the involvement of other TORC1-regulated signalling.

### *Mpk1 activation and* ADC17 *mRNA-actin patch association are necessary, but not sufficient, for Adc17 expression*

As CWI activation leads to TORC1 inhibition, we looked downstream of CWI activation to assess the sufficiency of *ADC17* mRNA relocalisation to actin patches and Mpk1 activation for Adc17 protein expression. We tethered *ADC17* mRNA to actin patches and used our inducible Mpk1 activation system (Figure S1D, 3A) (Williams et al., 2022). Combining these two features was insufficient to achieve Adc17 expression in the absence of rapamycin (Figure 3B). As our Mpk1 activation system requires the absence of glucose, we examined whether this was a factor by using *ede1Δ* cells with actin-patch tethered *ADC17* mRNA, as these cells have elevated Mpk1 activation (Figure S3F). These cells are likewise incapable of Adc17 expression in the absence of rapamycin. An additional factor(s) downstream of TORC1 inhibition must therefore also be involved.

**Figure 3:**
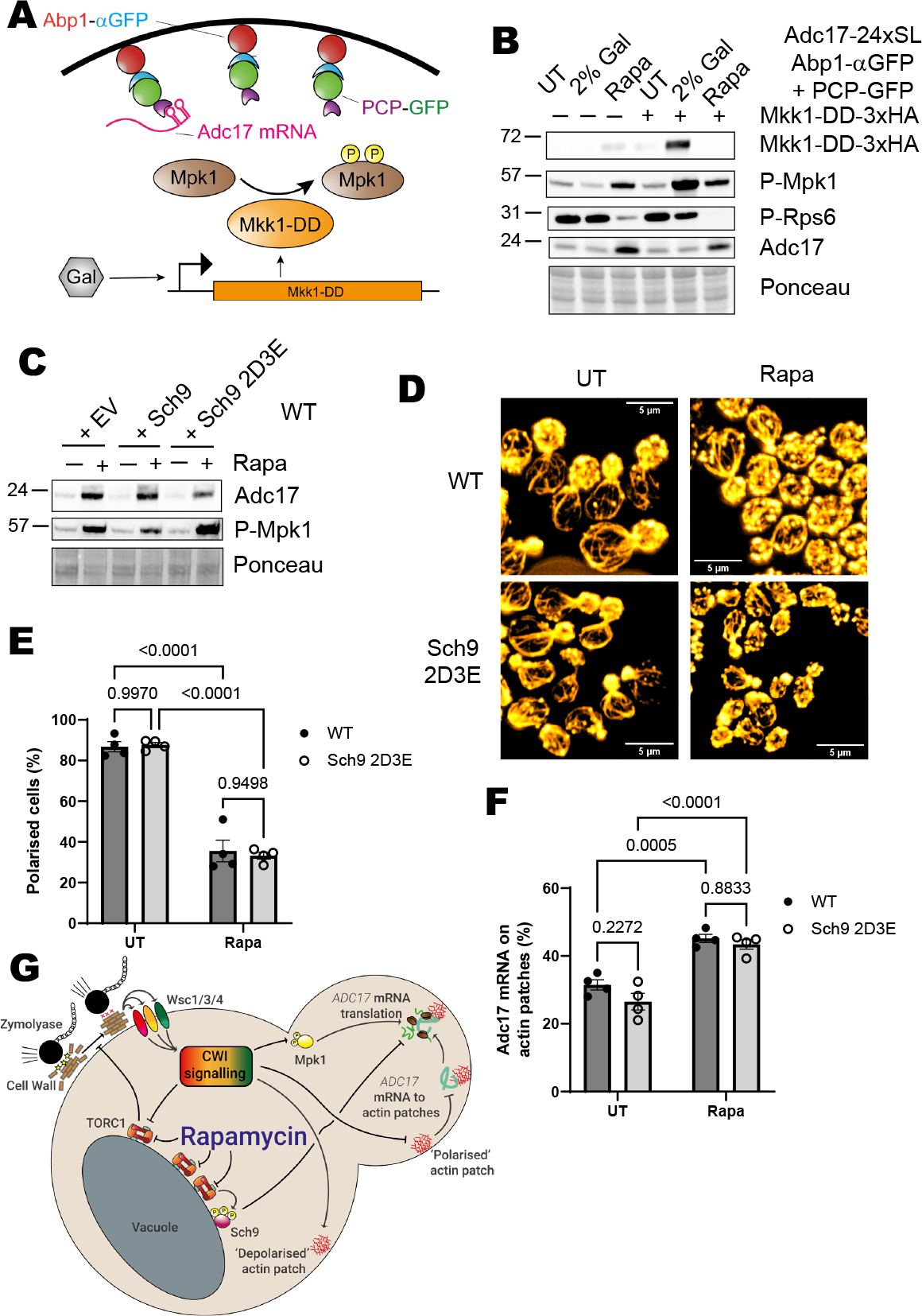
Sch9 inhibition is required for rapamycin-induced Adc17 expression by a CWI-independent mechanism. A) Scheme showing Mpk1 activation through inducible expression of the Mpk1 upstream kinase Mkk1 with the constitutively active ‘DD’ mutations and *ADC17* mRNA tethering to actin patches using the actin patch protein Abp1 tagged with an αGFP nanobody. B) Mkk1-DD-3xHA and Adc17 expression and Mpk1 and TORC1 activation (P-Mpk1 & P-Rps6 respectively) following 4 h rapamycin treatment or galactose induction in the cells with actin patch-tethered ADC17 mRNA (Adc17-24xSL Abp1-αGFP + PCP-GFP as described in Figure 3A. Representative of at least 3 experiments. C) Mpk1 activation (P-Mpk1) and Adc17 expression following 4 h rapamycin treatment in WT cells containing either empty vector (EV), WT Sch9, or Sch9-2D3E constitutively active mutant. Representative of at least 3 experiments. D) Effect of 1 h rapamycin treatment on actin polarisation in WT and Sch9-2D3E expressing cells. Representative of at least 3 experiments. Cellular actin is labelled with rhodamine-phalloidin. Scale bars show 5 µm. E) Quantification of images from D (n=4). F) Quantification of *ADC17* mRNAs localised to actin patches in WT and Sch9-2D3E expressing cells either untreated or treated with rapamycin for 1 h (n=4). G) Updated scheme from Figure 1A, showing additional role of Sch9 in Adc17 expression.

### Sch9 inhibition is important for Adc17 translation

To assess the involvement of factors downstream of TORC1, we returned to our mutant collection where signalling from individual TORC1 branches is maintained following TORC1 inhibition (Figure S2A). We assessed Adc17 expression in these mutants following rapamycin treatment, finding no change in autophagy-deficient and Pph2 deletion mutants, while Sit4 deletion had a small effect on Adc17 expression (Figures S3G-I). Excitingly, there was a notable decrease in Adc17 expression in Sch9-2D3E expressing cells, suggesting a role for Sch9 inhibition (Figure 3C).

We assessed whether Sch9 inhibition could modulate either Mpk1 activation or *ADC17* mRNA localisation. Sch9-2D3E expression did not compromise Mpk1 activation (Figure 3C), actin depolarisation (Figures 3D&E) or *ADC17* mRNA relocalisation to actin patches (Figure 3F). This validates the results from the previous section demonstrating the requirement for an additional factor to Mpk1 activation and *ADC17* mRNA relocalisation. TORC1 inhibition therefore promotes Adc17 expression through both CWI pathway activation and Sch9 inactivation (Figure 3G).

### Mpk1 and Sch9 are the major proteasome assembly pathway regulators

In a previous screen, we identified high levels of basal Adc17 expression in cells lacking the clathrin heavy chain, Chc1 (Williams et al., 2022). We examined whether the aspects of the proteasome assembly pathway we have thus far uncovered were active in *chc1Δ* cells. Under basal conditions, *chc1Δ* cells not only have elevated Adc17 expression, comparable to WT cells treated with rapamycin, they have extremely high levels of active Mpk1 and low levels of TORC1 activity, visualised by P-Rps6 (Figure 4A), consistent with our model (Figure 3G).

**Figure 4.**
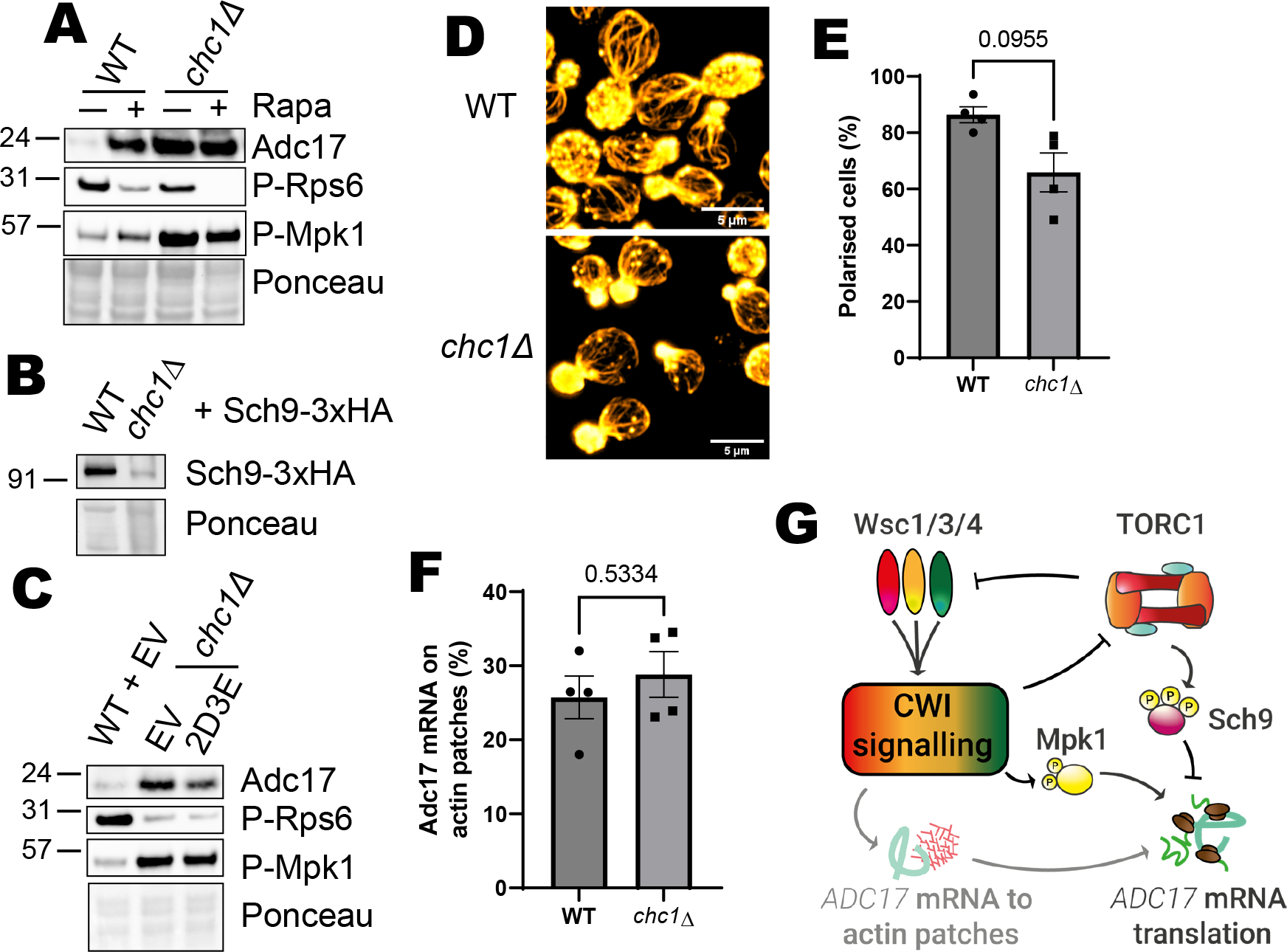
Absence of Chc1 chronically activates the proteasome assembly pathway through Mpk1 activation and Sch9 inhibition. A) Mpk1 activation (P-Mpk1) and Adc17 expression following rapamycin treatment in WT and *chc1Δ* cells. Representative of at least 3 experiments. B) Sch9 expression in WT and *chc1Δ* cells. Representative of at least 3 experiments. C) Mpk1 activity (P-Mpk1), TORC1 activity (P-Rps6) and Adc17 expression in WT and *chc1Δ* cells containing either empty vector (EV) or expressing Sch9-2D3E. Representative of at least 3 experiments. D) Actin polarisation in WT and *chc1Δ* cells. Representative of at least 3 experiments. Cellular actin is labelled with rhodamine-phalloidin. Scale bars show 5 µm. E) Quantification of images from D (n=4). F) Quantification of *ADC17* mRNAs localised to actin patches in untreated WT and *chc1Δ* cells (n=4). G) Scheme showing the signalling control of Adc17 expression.

We observed very low levels of Sch9 in *chc1Δ* cells, confirming that its overall function is decreased in *chc1Δ* cells compared to WT cells (Figure 4B). Increasing Sch9 activity by expressing the Sch9-2D3E phospho-mimetic protein reduced Adc17 expression without affecting the activation status of either TORC1 or Mpk1 (P-Rps6 and P-Mpk1, Figure 4C).

Surprisingly, given the low levels of TORC1 activity, *chc1Δ* cells largely maintain actin polarity (Figure 4D&E) (Henry et al., 2002). Consistently, a similar proportion of *ADC17* mRNA was localised to actin patches in WT and *chc1Δ* cells (Figure 4F). While *ADC17* mRNA relocalisation to actin patches may provide a boost to translation following TORC1 inhibition, it is therefore not necessary to sustain high levels of the protein when cells have chronic TORC1 inhibition and Mpk1 activation. It is important to note here that while there was not an increase in *ADC17* mRNA localised to actin patches, a significant proportion was still present. This level of localisation is likely to be sufficient to sustain the high level of protein when Mpk1 is active and Sch9 is inhibited.

Altogether, this work demonstrates that the proteasome assembly pathway is regulated by both Mpk1 and Sch9 kinases by TORC1 inhibition and CWI activation. CWI regulation of *ADC17* mRNA relocalisation to actin patches can provide a translational boost, but is not required under chronic TORC1 inhibition and CWI activation (Figure 4G).

## Supplementary Figure Legends

**Figure S1:**
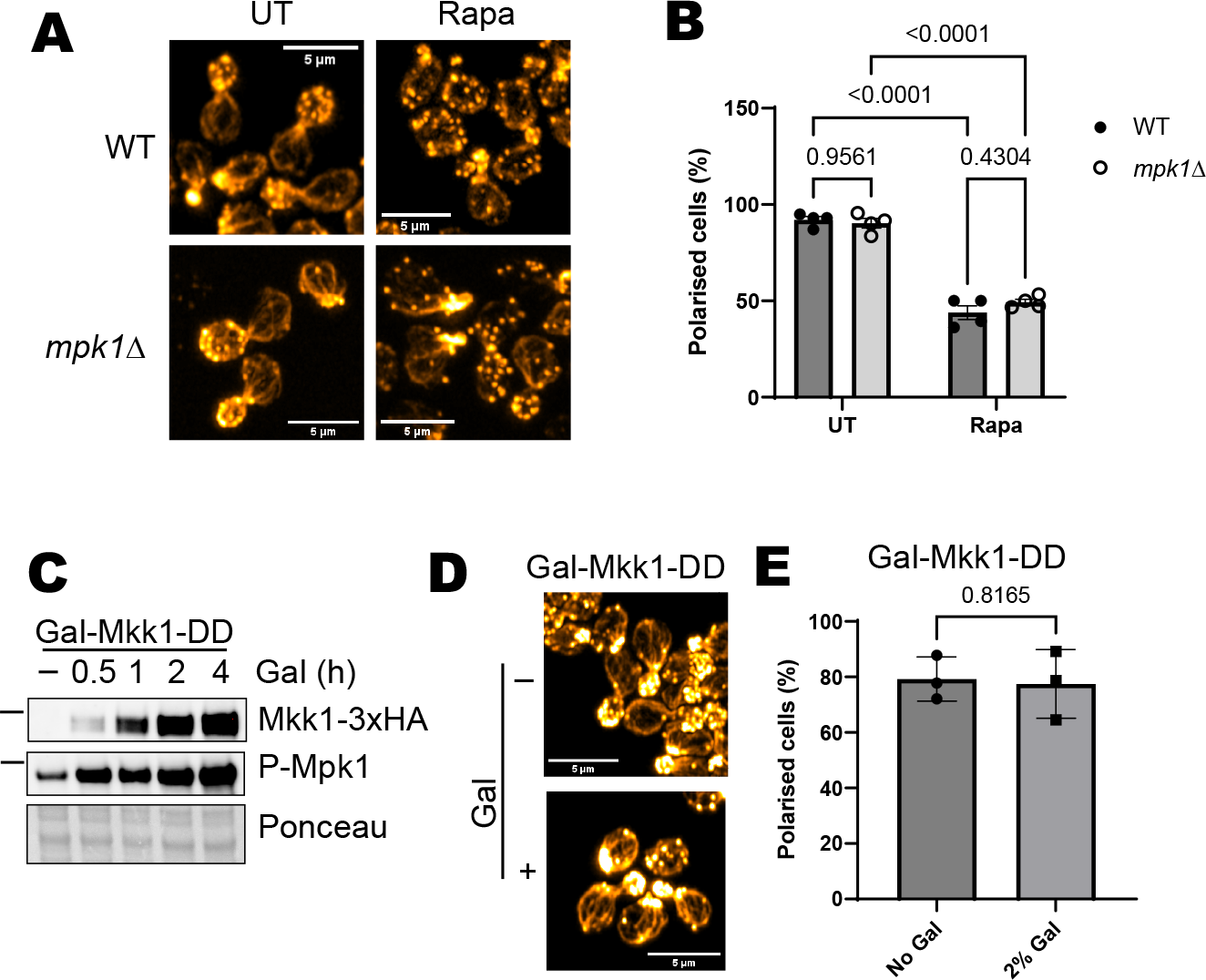
Related to Figure 1. Actin depolarisation following rapamycin treatment is activated by the CWI pathway independently of Mpk1 activation and Bni1 degradation. A) Effect of 1 h rapamycin treatment on actin polarisation in WT and *mpk1Δ* cells. Representative of at least 3 experiments. Cellular actin is labelled with rhodamine-phalloidin. Scale bars show 5 µm. B) Quantification of images from A (n=4). C) Induction of Mkk1-DD-3xHA from the Gal10 galactose inducible promoter, and consequent activation of Mpk1 (P-Mpk1) in WT cells. Representative of at least 3 experiments. D) Effect of Mkk1-DD-3XHA induction (1 h galactose) on actin polarisation in WT cells. Representative of at least 3 experiments. Cellular actin is labelled with rhodamine-phalloidin. Scale bars show 5 µm. E) Quantification of images from D (n=3).

**Figure S2:**
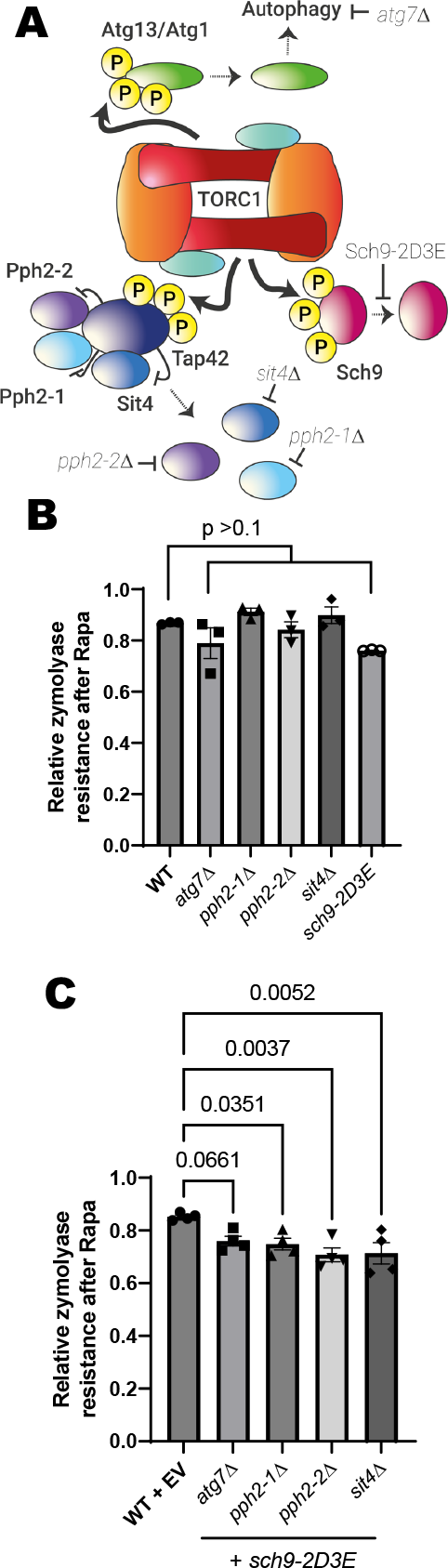
Related to Figure 2.TORC1 inhibts cell wall integrity activation through multiple pathways. A) TORC1 signalling to the autophagy, Tap42/PP2A phosphatase, and Sch9 branches. Mutants are displayed which prevent changes to each branch upon TORC1 inhibition. B) Resistance to 45 min zymolyase treatment after 1 h rapamycin treatment in the mutants indicated in Figure S2A. C) Resistance to 45 min zymolyase treatment after 1 h rapamycin treatment in the mutants indicated in Figure S2A expressing Sch9-2D3E.

**Figure S3:**
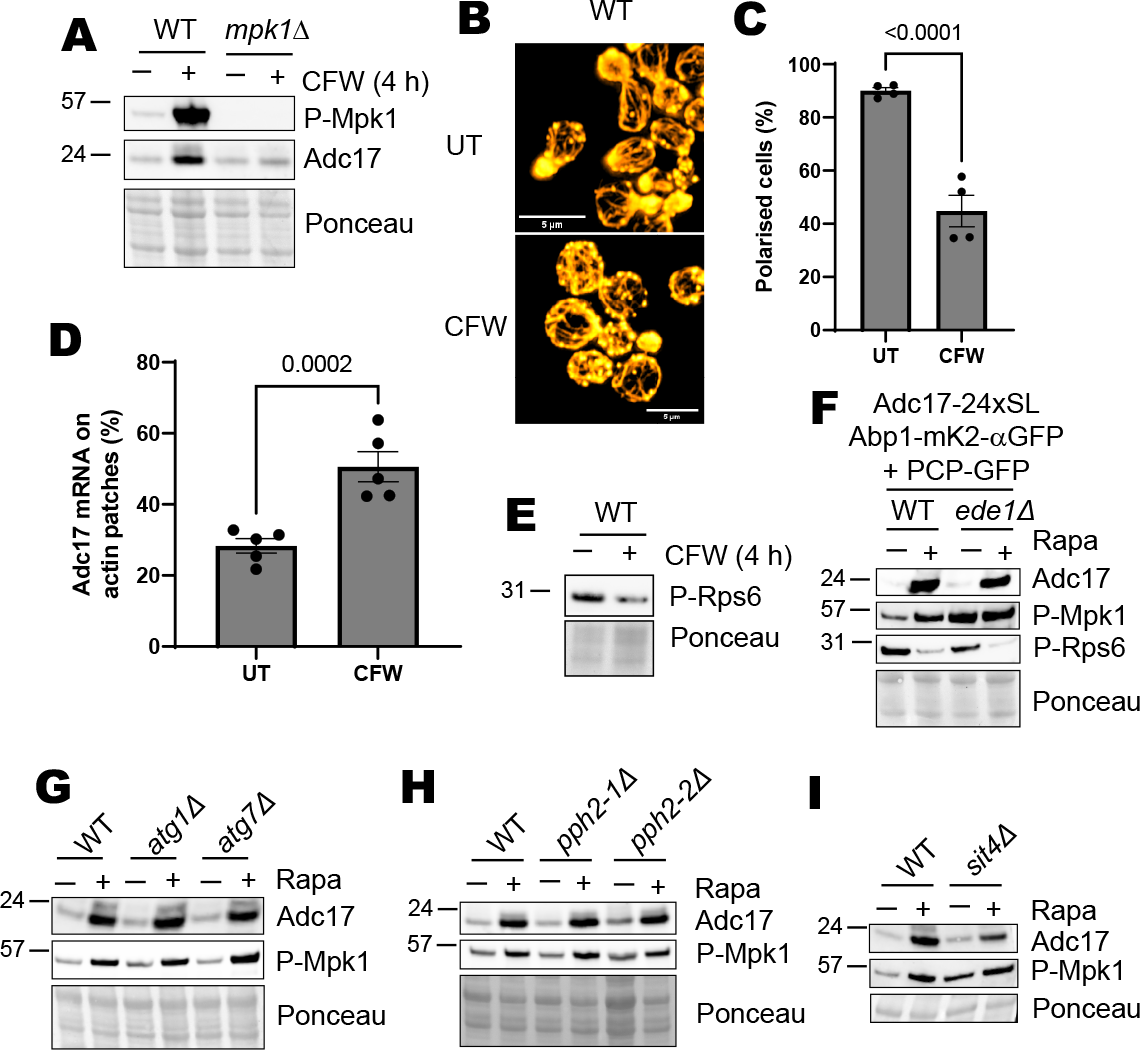
Related to Figure 3. Mpk1 activation and mRNA recruitment to cortical actin patches are insufficient to drive *ADC17* expression. A) Mpk1 activation (P-Mpk1) and Adc17 expression in WT and *mpk1Δ* cells after 4 h calcofluor white treatment. Representative of at least 3 experiments. B) Effect of 1 h calcofluor white treatment on actin polarisation in WT cells. Representative of at least 3 experiments. Cellular actin is labelled with rhodamine-phalloidin. Scale bars show 5 µm. C) Quantification of images from B (n=4). D) Quantification of *ADC17* mRNA localised to actin patches in untreated or 1 h calcofluor white treated WT cells (n=5). E) TORC1 activity (P-Rps6) in WT cells after 4 h calcofluor white treatment. Representative of at least 3 experiments. F) Adc17 expression and Mpk1 and TORC1 activity (P-Mpk1 & P-Rps6 respectively) untreated and following 4 h rapamycin treatment in WT and *ede1Δ* cells with *ADC17* mRNA tethered to actin patches. Representative of at least 3 experiments. G) Mpk1 activation (P-Mpk1) and Adc17 expression in WT and the autophagy deficient *atg1Δ* and *atg7Δ* mutants after 4 h rapamycin treatment. Representative of at least 3 experiments. H) Mpk1 activation (P-Mpk1) and Adc17 expression in WT, *pph2-1Δ*, and *pph2-2Δ* mutants after 4 h rapamycin treatment. Representative of at least 3 experiments. I) Mpk1 activation (P-Mpk1) and Adc17 expression in WT and *sit4Δ* cells after 4 h rapamycin treatment. Representative of at least 3 experiments.

**Table S1: Strains used in this article**

List of strains used in this article and information about their origin.

## Methods

### Yeast strains and growth/treatment conditions

All strains used were derived from the BY4741 collection and are listed in supplementary table 1. Mutants were made using homologous recombination as described previously (Agrotis et al., 2023; Black et al., 2023; Williams et al., 2022). Cells were grown at 30°C on YPD agar plates or in YEPD media, with shaking at 200 rpm. For all experiments cells were grown overnight prior to being used.

For drop assays, cells were adjusted to 0.2 OD_600 nm_ in YEPD media, and serially diluted 1/5 before being spotted on YEPD plates +/-20 ng/ml rapamycin (LC Laboratories, R5000) which were then imaged after 3 days.

For Western blot experiments, cell concentration was adjusted to 0.2 OD_600 nm_, and cells were grown until they reached ∼0.4 – 0.6 OD_600 nm_. They were then diluted back to 0.2 OD_600 nm_, split into 4 ml (untreated) and 12 ml (treated – 200 ng/ml rapamycin or 0.5 mg/ml calcofluor white) and returned to the incubator for 4 hours. Cells were then pelleted, flash-frozen in dry ice, and stored at -20°C overnight prior to extraction.

For galactose-inducible expression of Mkk1-DD from the Gal10 promoter to activate Mpk1, cells were grown to logarithmic phase in YEP + 2% Raffinose, before galactose was added to 2% to initiate expression.

For microscopy experiments, cells were split into 2 x 4 ml cultures and either fixed straight away with 3.7% formaldehyde (final concentration) or subjected to treatment (200 ng/ml rapamycin or 0.5 mg/ml calcofluor white (Sigma-Aldrich 910090)) for 1 hour, then fixed. Cells were incubated with the formaldehyde mixture for 20 minutes shaking at 30°C, then spun down and washed twice in PBS before actin was labelled using phalloidin and cells were mounted onto slides as described (Williams et al., 2022).

### Protein extraction and Western blot

Cell pellets were washed once in 600 ml ice-cold water, once in 400 ml ice-cold 2 M LiAc, and once in 400 ml ice-cold 0.4 M NaOH before resuspension in 110 ml lysis buffer containing protease and phosphatase inhibitors. This was placed in a heat block at 90°C for 10 minutes before addition of 2.64 ml 4 M acetic acid, and then returned to the heat block for 10 minutes. The supernatants were cleared by centrifugation, the protein concentration determined using a nanodrop, which was then normalised between samples.

Samples were diluted in 5x loading buffer and ∼30 mg run out on a 6-14% SDS-PAGE gel as described before semi-dry Western blotting (BioRad TurboID). Ponceau stained images were taken, membranes cut up and blocked with 5% milk for at least one hour, and primary antibodies diluted in TBS-T + 5% BSA + 0.02% NaOAc added (αP-Mpk1, Rabbit, 1:1000, CST, 4370S; αAdc17, Sheep, 1:200, DSTT, DU66321; αP-Rps6, Rabbit, 1:1000, CST, 2211; αHA, Mouse, 1:1000, Santa Cruz Sc-7392) overnight. In the morning, membranes were washed and incubated with appropriate secondary antibody diluted in TBS-T (HRP conjugated αMouse, 1:10000, CST, 7076S; HRP conjugated αRabbit, 1:10000, CST, 7074S; HRP conjugated αSheep, 1:5000, Sigma-Aldrich, A3415) for at least an hour before being washed and revealed (BioRad Clarity or Clarity Max ECL). All Western Blot imaging was performed using a BioRad Chemidoc system.

### Microscopy and analysis

Following sample preparation, cells were imaged on a Zeiss 880 using the Airyscan mode and a 63x lens to take Z-stacks covering the entire cell layer. These were subject to a Z-projection in FIJI, and actin polarisation and mRNA localisation were scored in FIJI as previously described (Williams et al., 2022). Statistical analysis was performed using 2-way ANOVA followed by multiple comparison t-tests in Graphpad PRISM.

### Zymolyase assays

5 ml of log phase cells, either treated with rapamycin for the indicated time or not, were spun down at 7200*g* for 3 minutes, and washed twice in 5 ml water. Cells were then resuspended in 1 ml 0.1M Tris (pH7.4), and the OD_600nm_ measured before and after 45 minutes incubation with 1.3 Units Zymolyase (Stratech).

